# On the transdiagnostic nature of peripheral biomarkers in major psychiatric disorders: a systematic review

**DOI:** 10.1101/086124

**Authors:** Jairo V. Pinto, Thiago C. Moulin, Olavo B. Amaral

## Abstract

The search for biomarkers has been one of the leading endeavors in biological psychiatry; nevertheless, in spite of hundreds of publications, it has yet to make an impact in clinical practice. To study how biomarker research has progressed over the years, we performed a systematic review of the literature to evaluate (a) the most studied peripheral molecular markers in major psychiatric disorders, (b) the main features of studies in which they are proposed as biomarkers and (c) whether their patterns of variation are similar across disorders. Out of the six molecules most commonly present as keywords in articles studying plasmatic markers of schizophrenia, major depressive disorder or bipolar disorder, five (BDNF, TNF-alpha, IL-6, C-reactive protein and cortisol) were the same across the three diagnoses. An analysis of the literature on these molecules showed that, while 65% of studies were cross-sectional and 66% compared biomarker levels between patients and controls in specific disorders, only 10% presented an objective measure of diagnostic or prognostic efficacy. Meta-analyses showed that variation in the levels of these molecules was robust across studies, but also similar among disorders, suggesting them to reflect transdiagnostic systemic consequences of psychiatric illness rather than diagnostic markers. Based on this, we discuss how current publication practices have led to research fragmentation across diagnoses, and what steps can be taken in order to increase clinical translation in the field.

## INTRODUCTION

With the growth of biological psychiatry and the widespread adoption of diagnostic classifications, the concept of diagnostic biomarkers has loomed promisingly in the horizon as a major goal of biological psychiatry (Kupfer *et al.* 2002; Insel & Quirion 2005). Although the idea of objective diagnostic tests in psychiatry is not new, and goes back to early promises such as the dexamethasone suppression test (Arana 1985), interest in biomarkers has grown exponentially over the last two decades, as shown by the steep rise in articles including the words “biomarker” and “psychiatry” (Fig. 1A). Not only part of the scientific literature, but also news pieces in scientific journals (Haag 2007) and in the lay media (Nutt 2016) have spread the idea that objective tests could eventually trump the centuries-old method of using clusters of symptoms to diagnose mental illness.

**Fig. 1.**
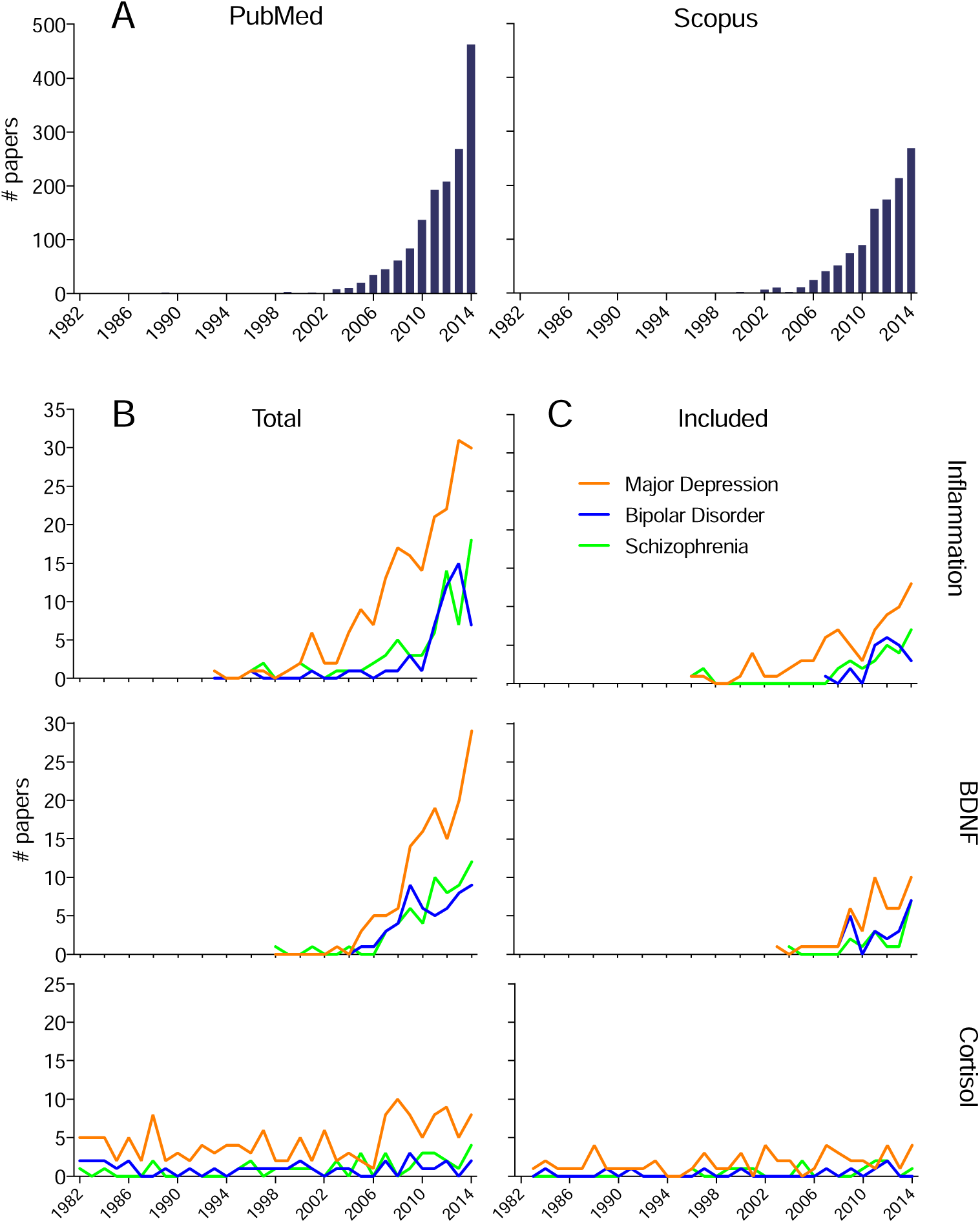
Growth of biomarker research in psychiatry (A) Number of PubMed (left) and Scopus (right) hits in searches for “biomarker” AND “psychiatry” or “psychiatric” for each year between 1982 and 2014 (B) Total number of articles over the same period retrieved for searches in PubMed and Scopus for “biomarker” AND either (a) “tumor necrosis factor alpha OR interleukin-6 OR C-reactive protein” (top) or (b) “BDNF” (middle) or (c) “hydrocortisone” (bottom) AND either (a) “major depression OR depressive disorder” (orange) (b) “bipolar disorder” (blue) or (c) “schizophrenia” (using Pubmed MeSH terms); (C) original articles fulfilling criteria for inclusion in our systematic review of experimental design features for each year.

Nevertheless, the application of biomarkers in the psychiatric clinic is still very limited. As recently discussed by various authors (Kapur *et al.* 2012; Lancet Psychiatry 2016), problems for translating research findings into clinical practice include the biological heterogeneity of diagnostic constructs (Heinrichs 2004; Hyman 2010; Pavão *et al.* 2015), the emphasis on statistical rather than clinical significance (Sterne & Davey Smith 2001; Perlis 2011), the use of extreme comparisons between prototypical patients and healthy controls (Kapur *et al.* 2012; Lancet Psychiatry 2016) and issues such as low statistical power, publication bias and lack of replication (Ioannidis 2005; Kapur *et al.* 2012). Moreover, the controversy over the soundness of the DSM as a framework for biological psychiatry (Cuthbert & Insel 2013), as well as evidence showing that major psychiatric disorders are promiscuous in terms of genetic loci (Smoller 2013), risk factors (Caspi *et al.* 2014) anatomical substrates (Goodkind *et al.* 2015; Sprooten *et al.* 2016) and treatment (Chouinard 2006), have led some to argue that the greatest promise in biomarker development might lie not in diagnosis, but in the prediction of prognosis and/or treatment response (Davis *et al.* 2014).

Although such advice seems generally sound, it is uncertain whether such concerns have made a significant impact on biomarker research. Moreover, due to the high degree of separation of psychiatric research into different disorders, the question of whether the promiscuity among disorders observed for genetics, anatomy and risk factors also applies to peripheral biomarkers has only been studied in isolated cases (Fernandes *et al.* 2013; Goldsmith *et al.* 2016). With this in mind, we decided to (a) systematically investigate what are the most studied plasmatic markers for major psychiatric disorders, (b) perform a systematic review of the experimental design features of articles evaluating them as biomarkers and (c) on the basis of available meta-analytical evidence, investigate whether variation in their levels occurs similarly across different diagnoses.

## METHODS

### Search strategy and biomarker selection

We initially searched the PubMed and Scopus databases on June 5^th^, 2015 for articles containing the terms “biomarker” OR “biomarkers” AND (“serum” OR “blood” OR “plasma” OR “plasmatic”) AND one of six psychiatric disorders: (a) “bipolar disorder”, (b) “major depression” OR “major depressive disorder”, (c) “schizophrenia”, (d) “post-traumatic stress disorder” OR “PTSD”, (e) “attention-deficit-hyperactivity disorder” OR “ADHD”, (f) “autism” OR “ASD” Although this search was non-exhaustive, as articles might study peripheral markers without using the term “biomarker”, our objective was to specifically select papers on this topic for an automated keyword search - thus, the main goal was to minimize the presence of literature not relating to the theme.

Based on search results, we used a MATLAB script (available upon request) to count the numbers of articles containing each individual term in the author and index keywords of the Scopus database (the current PubMed XML field for keywords, “Other Terms”, was not present before 2013 and thus could not be used). We then ranked words in order of frequency and manually scanned the tables to build a list of the molecules most frequently included as keywords for each individual disorder (Table 1), taking care to aggregate counts of terms relating to the same molecule (e.g. “BDNF” and “brain-derived neurotrophic factor”).

**Table 1.** Most frequent molecules among Scopus keywords in peripheral biomarker studies of different psychiatric disorders. Table shows the top-ranked endogenous molecules among Scopus keywords in a database search for “biomarker” AND “serum OR blood OR plasma OR plasmatic” AND individual disorder (see methods). Note that not all molecules included as keywords necessarily represent candidate peripheral biomarkers. The 5 top hits for mood disorders are in bold, and at least 3 of them appear in every disorder. Italic indicates molecules used as markers of treatment effects (e.g. prolactin, insulin) and molecules not used as peripheral markers, which appeared in the text for other reasons (e.g. glutamic acid, serotonin). ACTH: adenocorticotropic hormone; BDNF: brain-derived neurotrophic factor; CRH: corticotrophin releasing hormone; CRP: C-reactive protein; DOPAC: 3, 4 dihydroxyphenylacetic acid; EGF: epidermal growth factor; GABA: gamma-aminobutyric acid: IL-1 beta: interleukin 1 beta; IL-2: interleukin 2; IL-4: interleukin 4; IL-6: interleukin 6; IL-8: interleukin 8; IL-10: interleukin 10; MHPG: 4-hydroxy-3 methoxyphenylethylene glycol; NGF: nerve growth factor; S100B: S100 calcium-binding protein beta; SOD: superoxide dismutase; TNF-α: tumor necrosis factor alpha.

| MDD |  | BD |  | Schizophrenia |  | PTSD |  | Autism |  | ADHD |  |
| --- | --- | --- | --- | --- | --- | --- | --- | --- | --- | --- | --- |
| Molecule | # | Molecule | # | Molecule | # | Molecule | # | Molecule | # | Molecule | # |
| **BDNF** | **86** | **BDNF** | **55** | **BDNF** | **48** | **Cortisol** | **19** | Glutathione | 20 | *Dopamine* | 7 |
| **Cortisol** | **61** | **TNF-α** | **21** | *Prolactin* | *31* | **CRP** | **12** | Serotonin | 20 | Norepinephrine | 5 |
| **IL-6** | **60** | **IL-6** | **18** | **TNF-α** | **22** | **IL-6** | **11** | **BDNF** | **12** | **BDNF** | **4** |
| **CRP** | **53** | **Cortisol** | **13** | **IL-6** | **19** | *CRH* | 7 | Homocysteine | 12 | Adiponectin | 4 |
| **TNF-α** | **46** | **CRP** | **12** | Glutathione | 17 | **BDNF** | **6** | **TNF-α** | **10** | **Cortisol** | **3** |
| Fibrinogen | 23 | S100B | 11 | *Insulin* | *17* | ACTH | 6 | **IL-6** | **9** | *Oxyhemoglobin* | 3 |
| IL-1 beta | 21 | SOD | 10 | *Leptin* | *15* | IL-8 | 5 | Cysteine | 8 | *Serotonin* | 3 |
| *Serotonin* | *18* | *Glutamic acid* | *10* | *Glucose* | *14* | Neuropeptide Y | 5 | Interferon-γ | 8 | *Cholesterol* | 2 |
| IL-10 | 16 | IL-10 | 8 | **Cortisol** | **13** | IL-1 beta | 4 | Oxytocin | 8 | **IL-6** | **2** |
| S100B | 16 | Neurotrophin-3 | 8 | *Dopamine* | 13 | IL-4 | 4 | Lactic acid | 7 | **CRP** | **2** |
| *Glutamic acid* | *13* | NGF | 7 | **CRP** | **12** | Interferon-γ | 4 | Methionine | 7 | MHPG | 2 |
| *Prolactin* | *12* | GABA | 7 | GABA | *12* | **TNF-α** | **4** | Melatonin | 7 | Ferritin | 2 |
| *Insulin* | *11* | IL-1 beta | 6 | ACTH | 12 | Annexin a2 | 4 | *Glutamic acid* | 7 | Homovanillic acid | 2 |
| GABA | *11* | IL-2 | 6 | Interferon-γ | 11 | Serotonin | 4 | GABA | 6 | Neuropeptide Y | 2 |
| Interferon-γ | 10 | Nitric oxide | 6 | Homocysteine | 11 | S100B | 4 | IL-1 beta | 5 | DOPAC | 2 |
| Kynurenine | 10 | *Dopamine* | 6 | IL-10 | 11 | IL-2 | 3 | EGF | 5 | *Iron blood level* | 2 |
| *Leptin* | *10* | *Insulin* | 6 | *Glutamic acid* | *11* | *Leptin* | 3 | *Various* | - | *Various* | - |

### Selection criteria and analysis of original article features

Using the three disorders with the largest number of articles in our search (major depressive disorder, schizophrenia and bipolar disorder), we chose five molecules appearing among the top six keywords for these disorders - brain-derived neurotrophic factor (BDNF), tumor necrosis factor alpha (TNF-α), interleukin-6 (IL-6), C-reactive protein (CRP) and hydrocortisone (cortisol) for further study. We then performed searches in PubMed and Scopus on August 31^st^, 2015 using the following terms as keywords (as well as MeSH terms for all of them in PubMed):

(a) one of the three disorders (“schizophrenia” / “bipolar disorder” / “major depression” OR “major depressive disorder”)
(b) one of the biomarkers above, with inflammatory markers being pooled into a single search due to the high crossover of articles among them (“BDNF” / “interleukin-6” OR “c-reactive protein” OR “tumor necrosis factor alpha” / “hydrocortisone”).
(c) “biomarker” OR “biomarkers” For each biomarker/disorder combination, we screened abstracts (or full text when necessary) for the following inclusion criteria: (a) original articles, (b) in English, (c) including human patients (d) with the disorder of interest, according to DSM or ICD criteria and (e) performing peripheral measurements of the molecule in question (for a summary of the search strategy, see Fig. 2A). We focused on peripheral levels of the molecules, and did not include articles evaluating genetic polymorphisms and/or epigenetic changes in their corresponding genes. We then obtained the full text of every article (except for five articles for which it could not be obtained) and extracted the following information on experimental design features (presented in Table 2, Fig. 3, Supplementary Tables 1-3 and Supplementary Fig. 1):

(a) the specific molecule measured (e.g. BDNF protein, proBDNF protein, BDNF mRNA), site of measurement (e.g. blood, saliva, sweat, urine, hair) and any specific conditions of measurement (e.g. after dexamethasone challenge).
(b) the group comparisons performed in the article (e.g. patients vs. controls; different states of the disorder; patients with the disorder vs. those with another disorder; before vs. after treatment).
(c) the correlations performed within the group of patients (e.g. with symptoms, with illness progression, with treatment response or prognosis, with other markers).
(d) whether the study was cross-sectional or longitudinal.
(e) for longitudinal studies, whether it belonged to a predictive study in a population without the disorder.
(f) whether it offered an objective measure of diagnostic/prognostic efficacy (e.g. receiver operating characteristic (ROC) curves, odds ratio, classifier accuracy) or only presented a statistical comparison between groups.
(g) the total number of markers studied in the article.
(h) the total sample size of the article.
(i) the journal in which it was published.

Information was extracted by two of the authors (O.B.A. and J.V.P.) after extensive discussion of criteria, and 20% of the articles were cross-checked by both investigators, yielding a kappa coefficient of .948 for categorical variables. Controversies were solved with the participation of a third investigator (T.C.M.).

**Fig. 2.**
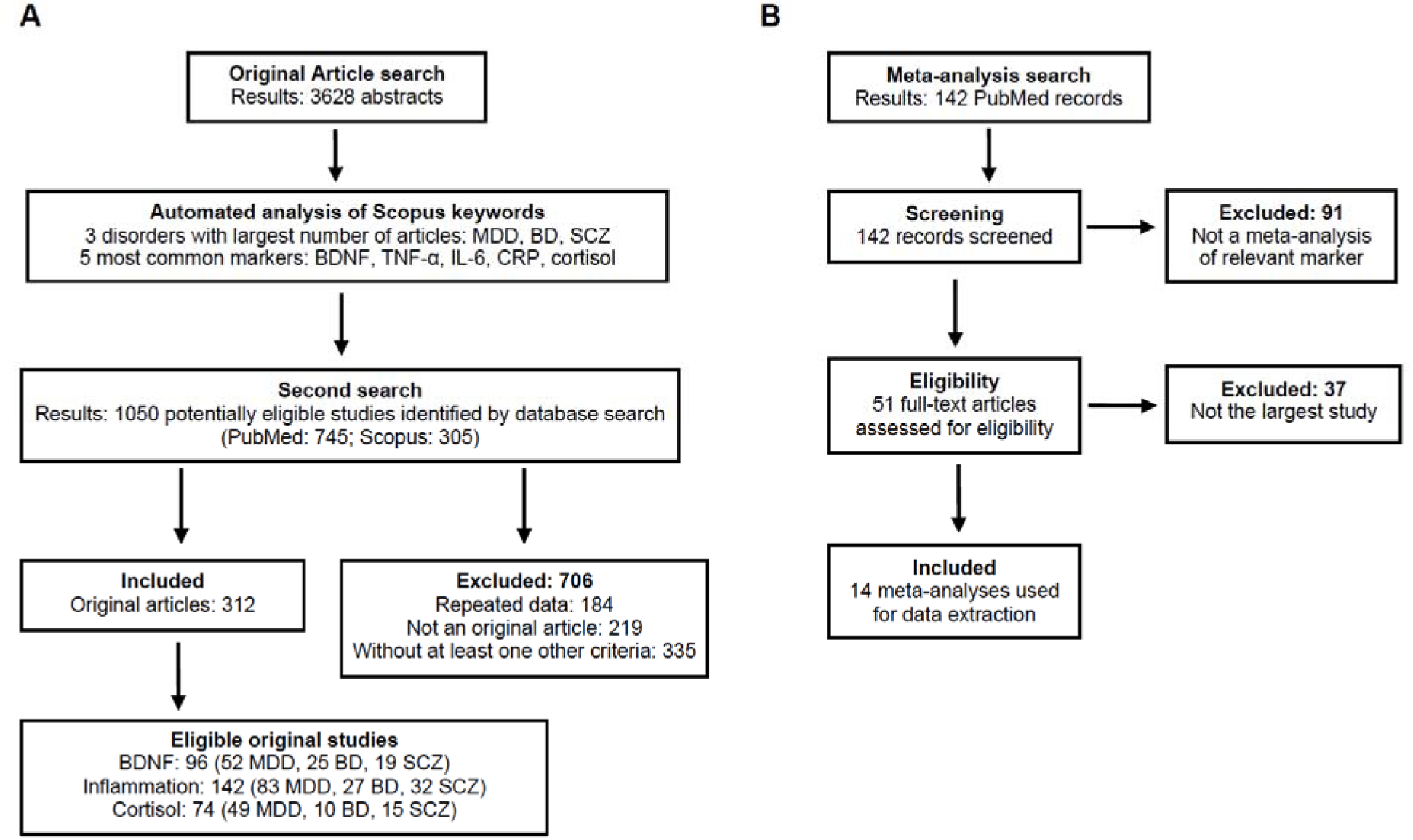
Flowchart depicting the selection of studies. Flowchart detailing the various stages of study selection in the studies. (A) Search strategy for original articles. Eligible studies for each biomarker include articles appearing in more than one search – thus, the sum of articles for individual disorders is greater than the total number of articles for each marker. **(B)** Search strategy for meta-analyses. A more detailed account of the search procedures can be found in the methods section and in the Supplementary Appendix. MDD, major depressive disorder; BD, bipolar disorder; SCZ, schizophrenia; BDNF, brain-derived neurotrophic factor; TNF-α, tumor necrosis factor alpha; IL-6, interleukin-6, CRP, C-reactive protein.

**Figure 3.**
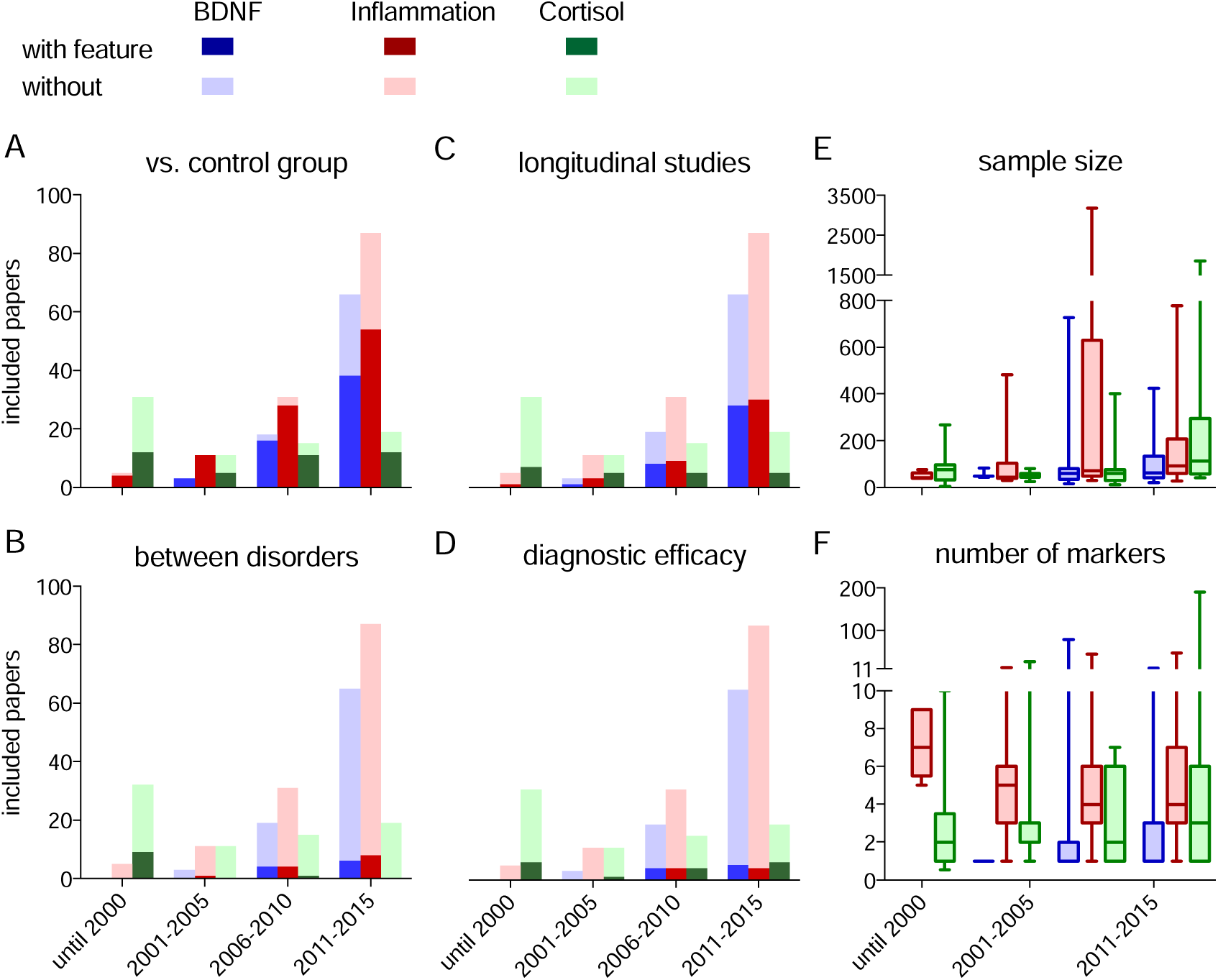
Variations of experimental design features of articles over different periods. Figure shows the frequency of **(A)** patient vs. control comparisons, **(B)** comparisons between disorders, **(C)** comparisons including a measure of diagnostic efficacy and **(D)** longitudinal studies, as well as the distribution of **(E)** sample size and **(F)** number of peripheral biomarkers studied among analyzed articles measuring BDNF (blue), TNF-α/IL-6/CRP (red) and cortisol (green) over 4 epochs (until 2000, 2001-2005, 2006-2010, 2011- 2015). For dichotomous variables (A-D), bars represent the total number of articles, with dark shading representing articles with each experimental design feature and light shading indicating those without them. Distributions of quantitative variables (E-F) are expressed as box-whisker plots (center line, median; box, interquartile range, whiskers, 5^th^/95^th^ percentiles). Spearman’s ρ coefficients for correlations between year of publication and each feature are (A) ρ=−0.01, p=0.80, (B) ρ=−0.13, p=0.025, (C) ρ=−0.04, p=0.47, (D) ρ=0.08, p=0.20, (E) ρ=0.21, p=4x10^−4^, (F) ρ=−0.003, p=0.95. Variations of additional features can be visualized in Supplementary Fig. 1.

**Table 2.**
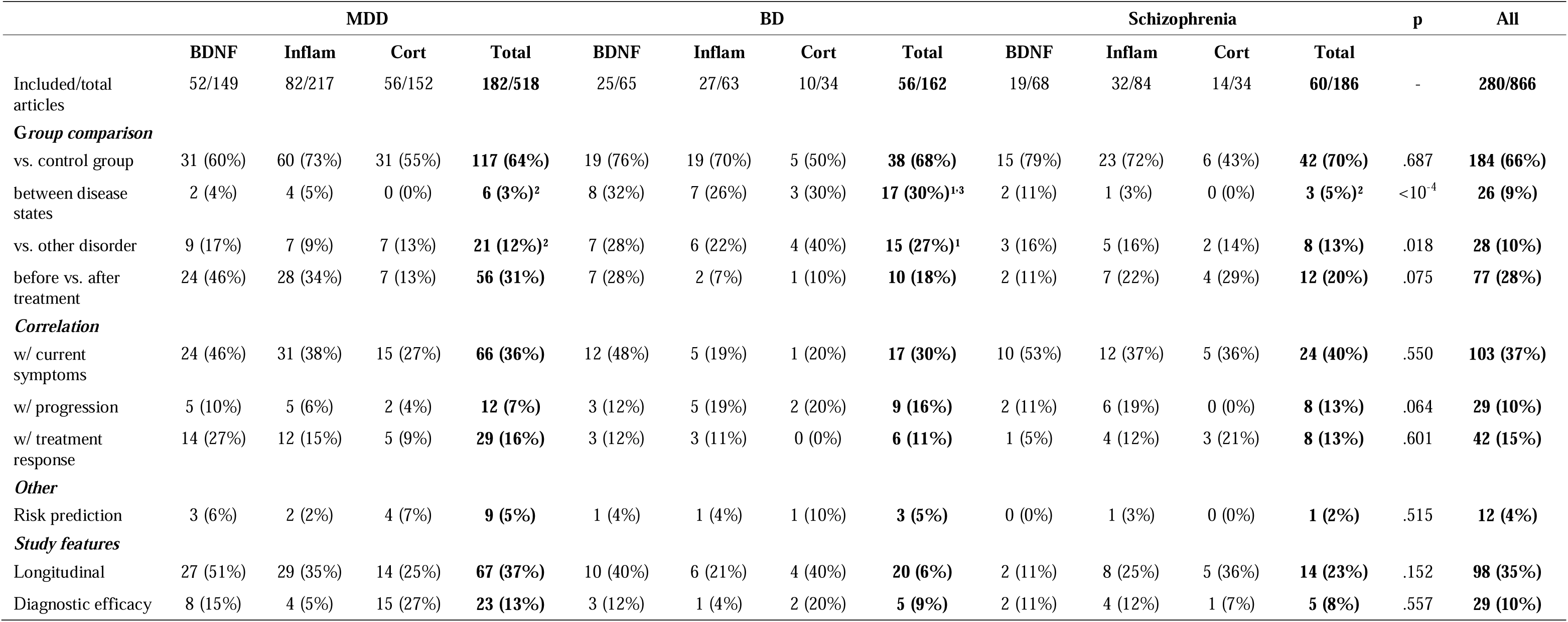
Experimental design features of retrieved articles for each biomarker/disorder combination. Articles measuring more than one marker for a given disorder or the same marker for more than one disorder were counted only once to calculate totals. BDNF, brain-derived neurotrophic factor; MDD, major depressive disorder; BD: bipolar disorder; SCZ: schizophrenia; p values refer to a χ^2^ test comparing aggregate values for all markers between the 3 disorders ^1^p<0.05 vs. MDD, ^2^p<0.05 vs. BD, ^3^ p<0.05 vs. SCZ, Fisher’s exact test comparing pairs of disorders. A similar table with aggregated totals for individual markers instead of disorders is presented as Supplementary Table 2.

|  | MDD |  |  |  | BD |  |  |  | Schizophrenia |  |  |  | p | All |
| --- | --- | --- | --- | --- | --- | --- | --- | --- | --- | --- | --- | --- | --- | --- |
|  | BDNF | Inflam | Cort | Total | BDNF | Inflam | Cort | Total | BDNF | Inflam | Cort | Total |  |  |
| Included/total articles | 52/149 | 82/217 | 56/152 | **182/518** | 25/65 | 27/63 | 10/34 | **56/162** | 19/68 | 32/84 | 14/34 | **60/186** | - | **280/866** |
| **Group comparison** |  |  |  |  |  |  |  |  |  |  |  |  |  |  |
| vs. control group | 31 (60%) | 60 (73%) | 31 (55%) | **117 (64%)** | 19 (76%) | 19 (70%) | 5 (50%) | **38 (68%)** | 15 (79%) | 23 (72%) | 6 (43%) | **42 (70%)** | .687 | **184 (66%)** |
| between disease states | 2 (4%) | 4 (5%) | 0 (0%) | **6 (3%)<sup>2</sup>** | 8 (32%) | 7 (26%) | 3 (30%) | **17 (30%)<sup>1,3</sup>** | 2 (11%) | 1 (3%) | 0 (0%) | **3 (5%)<sup>2</sup>** | <10<sup>-4</sup> | **26 (9%)** |
| vs. other disorder | 9 (17%) | 7 (9%) | 7 (13%) | **21 (12%)<sup>2</sup>** | 7 (28%) | 6 (22%) | 4 (40%) | **15 (27%)<sup>1</sup>** | 3 (16%) | 5 (16%) | 2 (14%) | **8 (13%)** | .018 | **28 (10%)** |
| before vs. after treatment | 24 (46%) | 28 (34%) | 7 (13%) | **56 (31%)** | 7 (28%) | 2 (7%) | 1 (10%) | **10 (18%)** | 2 (11%) | 7 (22%) | 4 (29%) | **12 (20%)** | .075 | **77 (28%)** |
| **Correlation** |  |  |  |  |  |  |  |  |  |  |  |  |  |  |
| w/ current symptoms | 24 (46%) | 31 (38%) | 15 (27%) | **66 (36%)** | 12 (48%) | 5 (19%) | 1 (20%) | **17 (30%)** | 10 (53%) | 12 (37%) | 5 (36%) | **24 (40%)** | .550 | **103 (37%)** |
| w/ progression | 5 (10%) | 5 (6%) | 2 (4%) | **12 (7%)** | 3 (12%) | 5 (19%) | 2 (20%) | **9 (16%)** | 2 (11%) | 6 (19%) | 0 (0%) | **8 (13%)** | .064 | **29 (10%)** |
| w/ treatment response | 14 (27%) | 12 (15%) | 5 (9%) | **29 (16%)** | 3 (12%) | 3 (11%) | 0 (0%) | **6 (11%)** | 1 (5%) | 4 (12%) | 3 (21%) | **8 (13%)** | .601 | **42 (15%)** |
| **Other** |  |  |  |  |  |  |  |  |  |  |  |  |  |  |
| Risk prediction | 3 (6%) | 2 (2%) | 4 (7%) | **9 (5%)** | 1 (4%) | 1 (4%) | 1 (10%) | **3 (5%)** | 0 (0%) | 1 (3%) | 0 (0%) | **1 (2%)** | .515 | **12 (4%)** |
| **Study features** |  |  |  |  |  |  |  |  |  |  |  |  |  |  |
| Longitudinal | 27 (51%) | 29 (35%) | 14 (25%) | **67 (37%)** | 10 (40%) | 6 (21%) | 4 (40%) | **20 (6%)** | 2 (11%) | 8 (25%) | 5 (36%) | **14 (23%)** | .152 | **98 (35%)** |
| Diagnostic efficacy | 8 (15%) | 4 (5%) | 15 (27%) | **23 (13%)** | 3 (12%) | 1 (4%) | 2 (20%) | **5 (9%)** | 2 (11%) | 4 (12%) | 1 (7%) | **5 (8%)** | .557 | **29 (10%)** |

Percentages of articles with or without each experimental design feature were initially calculated for each disorder/marker combination. We then calculated aggregate percentages for each disorder or marker and for the whole sample of articles; in these cases, articles appearing in more than one search were counted only once. Articles in which a particular feature was present for one marker/disorder but not for other(s) were considered to include that feature when calculating aggregate percentages. The complete database of articles retrieved by the search, as well as their categorization for each experimental design feature, is available in tables presented as Supplementary Data.

### Meta-analysis search and biomarker evaluation across disorders

In order to study whether variation in biomarker levels is similar or distinct among the three analyzed disorders, we searched PubMed on December 1^st^, 2016 for articles containing the terms (“meta-analysis”[Publication Type] OR “meta-analysis as topic” [MeSH Terms] OR “meta-analysis”[All Fields]) AND (schizophrenia OR bipolar disorder OR major depression OR major depressive disorder) AND (brain-derived neurotrophic factor OR bdnf OR interleukin-6 OR il-6 OR c-reactive protein OR tumor necrosis factor-alpha OR tnf-alpha OR cortisol OR hydrocortisone), as well as corresponding MeSH terms.

We included meta-analyses studying levels of at least one of the five biomarkers (i.e. BDNF, IL-6, TNF-alpha, C-reactive protein, cortisol) in at least one of the three studied disorders (i.e. MDD, bipolar disorder, schizophrenia) in which patient vs. control comparisons were performed (including comparisons for specific disease states). When more than one meta-analysis was available for the same biomarker, disorder and state, we selected the one with the largest sample size for inclusion in Table 3. Titles and abstracts were screened independently by two of the authors (J.V.P. and T.C.M.), and in case of disagreement consensus was reached using a third investigator (O.B.A). The search process for meta-analyses is described on Fig. 2B.

**Table 3.**
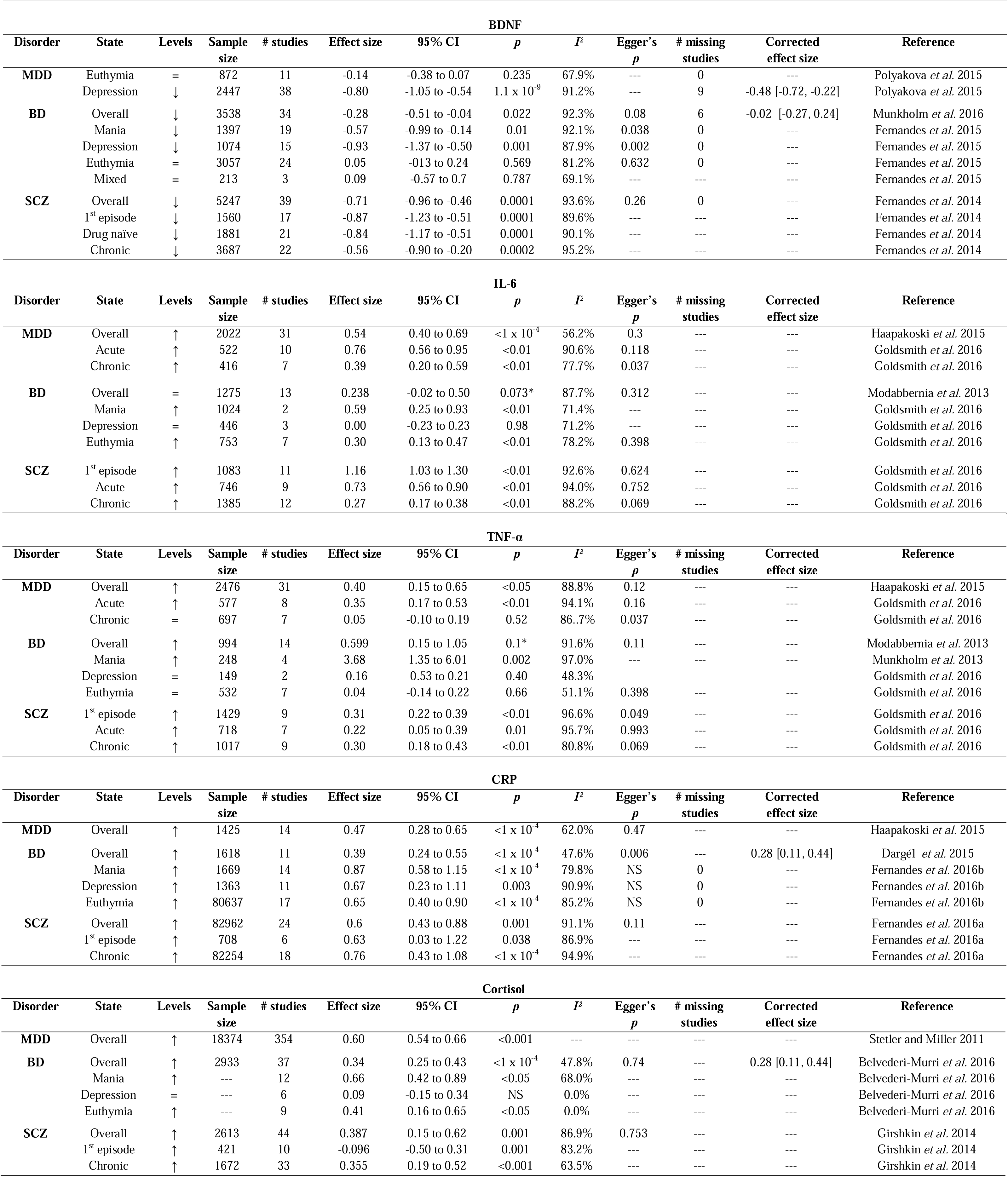
Meta-analytic estimates of variation in serum levels of different markers across disorders. The table describes the variation in serum/plasma levels of each marker for patient vs. control comparisons in each disorder, as well as in different states of the disorder when available. For each comparison, we used the available meta-analysis with the largest sample size in our search (see Methods). Columns show the direction of variation, total sample size and number of studies included in the meta-analysis, effect size with 95% confidence interval, *p* value for the comparison, heterogeneity measured by I^2^, *p* value for publication bias analyses (usually performed by Egger’s test), number of missing studies estimated through funnel plot/trim-and-fill analysis, corrected effect size after the imputation of missing studies and reference. State comparisons were defined according to the original meta-analysis. Effect sizes are given as provided in the original meta-analyses (Cohen’s d, Hedges’ g or standardised mean deviation). In cortisol meta-analyses, the overall comparison for depression used serum, salivary and urinary cortisol measured either at basal levels or after pharmacological challenges; the overall comparison for bipolar disorder used basal serum or salivary levels measured at any time; state comparisons in bipolar disorder and all comparisons in schizophrenia used morning serum or salivary cortisol levels only. --- stands for absence of information in the included article. * indicates p values corrected for multiple comparisons in the original meta-analysis. MDD; major depressive disorder; BD, bipolar disorder; SCZ, Schizophrenia; BDNF, brain-derived neurotrophic factor; IL-6, interleukin 6; TNF-alpha, tumor necrosis factor alpha.

For each meta-analysis, we extracted sample size, number of studies, effect size with 95% confidence intervals, p value and the I^2^ metric as a measure of heterogeneity. We also extracted *p* values for publication bias tests, the number of missing studies when funnel plot/trim-and-fill analyses were performed, and the corrected effect size after accounting for these studies when these were available. These data are included in Table 3 as well as in Supplementary Tables 4 to 8.

### Statistical analysis

For comparisons of categorical experimental design features between articles on different markers or different disorders, we used chi-square tests for omnibus comparisons followed by Fisher’s exact test between specific pairs of markers/disorders. For comparisons of quantitative features (i.e. sample size and number of markers) between articles on different markers or disorders, we used Kruskal-Wallis tests with Dunn’s test as a post-hoc. For correlations of experimental design features of articles with the year of publication, we used rank-biserial correlations for categorical variables and Spearman’s nonparametric correlations for quantitative variables (as both year and sample size/number of markers presented heavily skewed distributions).

## RESULTS

Our automated keyword search (see flowchart in Fig. 2) revealed that, out of the six molecules most commonly present as keywords in articles retrieved using “biomarker” and schizophrenia, major depressive disorder or bipolar disorder, five (BDNF, TNF-alpha, IL-6, C-reactive protein and cortisol) were the same across these disorders (Table 1). The temporal distribution of these articles (as well as of those fulfilling inclusion criteria) is displayed on Fig. 1 (B and C), revealing that articles on BDNF and inflammatory markers increased sharply after around 2005, whereas the number of articles on cortisol has remained relatively stable over two decades. Most articles on BDNF and inflammatory markers studied serum levels of these proteins, while cortisol articles usually studied serum or salivary cortisol levels, with frequent use of pharmacological challenges such as the dexamethasone suppression test as well (Supplementary Table 1).

An analysis of experimental design features of the literature on these molecules (Table 2 and Supplementary Table 2) showed that, whilst 66% of articles performed comparisons between patients and healthy controls, only 35% were longitudinal studies. Moreover, only 10% presented an objective measure of diagnostic or prognostic efficacy, such as a receiver operating characteristic (ROC curve) or an odds ratio for values above a certain threshold (as detailed in Supplementary Table 3). Most of these numbers did not vary significantly across markers or disorders, but some differences were observed: (a) state comparisons and comparisons with other disorders were more frequent for bipolar disorder, (b) pre vs. post-treatment comparisons and correlations of levels with symptoms were more common for BDNF and less frequent for cortisol and (c) cortisol studies presented patient vs. control comparisons less frequently, but included measures of diagnostic efficacy much more commonly than those on other markers (especially inflammation). Median sample size was 71.5 (interquartile range, 45.2-140.8) and larger for inflammatory markers than for BDNF or cortisol (p=0.004, Kruskal-Wallis test). The median number of markers addressed in each study was 3 (interquartile range, 1-5), and also larger for studies on inflammation (p<10^−4^, Kruskal-Wallis test).

Temporal trends for various types of comparison, as well as for median sample size and number of markers, are shown on Fig. 3 and Supplementary Fig. 1. No major differences were observed for the frequency of specific comparisons or correlations over time, and rank-biserial correlations testing for a time-related trend did not yield p values under 0.05 except for a slight decrease in the frequency of between-disorder comparisons (ρ=−0.13, p=0.025). Nevertheless, increases in the total number of articles meant that absolute numbers of articles tended to increase for most categories. Sample size also tended to increase over time (addressed in each study was =0.21, p=4×10^−4^), although its distribution varied widely in all periods.

We then moved on to analyze the variation of the five markers in each of the three disorders. Table 3 shows the results of the largest meta-analysis of patient vs. control comparisons for each marker/disorder combination (in various states of the disorder when available) (Stetler & Miller 2011; Munkholm *et al.* 2013, 2015; Modabbernia *et al.* 2013; Fernandes *et al.* 2014, 2015, 2016a, 2016b; Girshkin *et al.* 2014; Dargel *et al.* 2015; Polyakova *et al.* 2015; Haapakoski *et al.* 2015; Belvederi Murri *et al.* 2016; Goldsmith *et al.* 2016). Statistically significant variation in the level of all five molecules was found to occur in at least some states of all three disorders. Interestingly, the direction of variation was invariably the same across disorders for all markers (i.e. reductions in BDNF, increases in inflammatory markers and cortisol).

In mood disorders, differences between patients and controls were more marked among acutely ill patients (especially for mania in bipolar disorder) in the case of BDNF, IL-6 and TNF-α, while alterations in schizophrenia were observed both in acute and chronic illness for all markers. Cortisol and C-reactive protein levels appeared to be more fixed across disease states in mood disorders, with variations observed in euthymia as well. Effect sizes were generally comparable across disorders; however, evidence was more robust in some cases. Meta-analyses of BDNF, cortisol and C-reactive protein tended to yield more reliable differences than those studying IL-6 and TNF-α for mood disorders, while in schizophrenia differences were robust for all molecules. Heterogeneity between studies was almost invariably large, and evidence for publication bias was observed in a reasonable number of cases, although corrected effect sizes were infrequently provided.

For analyses other than patient vs. control comparisons (e.g. correlations with symptoms, illness progression, prediction of treatment response, etc.), meta-analytic results were not always available; moreover, results of individual studies were frequently contradictory. This might be related to differences in the clinical samples studied, but also to the low reliability of these secondary analyses in most articles (see Discussion). We summarized results for these additional analyses in Supplementary Tables 4-8, using meta-analyses whenever possible, or individual studies from our sample when meta-analyses were not available. One should note, however, that our search strategy for original studies, unlike the one used for metaanalyses, was not planned to be exhaustive (due to use of the term “biomarker”); thus, these tables should not be taken to include the full range of available literature on the subject.

## DISCUSSION

Although the promiscuity of peripheral biomarkers across diagnoses has received relatively little systematic attention in the literature, an automated keyword search revealed that the most frequently studied peripheral biomarkers are generally the same across major psychiatric disorders. Besides the biomarkers we chose to focus on (BDNF, IL-6, TNF-α, CRP and cortisol), other molecules were also studied in many disorders, such as oxidative stress markers (glutathione), other cytokines (IL-10, IL-1β) and the astrocytic protein S100B (Table 1). This is not in itself surprising, as one might expect that literature trends will promote interest in similar molecules across disorders. However, the fact that variation patterns were also similar across diagnoses suggests that there are real biological commonalities among psychiatric disorders in terms of their peripheral manifestations.

The reason for this similarity is worthy of discussion. On one hand, it can be thought of as a sign of overlap between the pathophysiology of major psychiatric disorders, a fact suggested by genetic (Smoller 2013), risk factor (Caspi *et al.* 2014) and neuroanatomical (Goodkind *et al.* 2015; Sprooten *et al.* 2016) similarities among diagnoses. After all, symptom-defined diagnostic boundaries in psychiatry should not be expected to “carve nature at its joints” all the way to the molecular level (Kendler 2006; Pavão *et al.* 2015). An additional interesting finding, though, is that many of the markers in question, such as cytokines and cortisol, are increased by various forms of chronic stress in both animal models and humans (McEwen 2007). This suggests that they might be related to the general “wear and tear” or allostatic load associated with mental or clinical illness (Kapczinski *et al.* 2008), rather than to the specific pathophysiology of any given disorder.

Also in favor of this view is the fact that alterations in cytokines and cortisol can be observed in acute stress models even in normal volunteers (Dickerson & Kemeny 2004; Steptoe *et al.* 2007; Slavich *et al.* 2010). And although a BDNF response to acute stress in humans has not been shown, there is evidence that its levels may be altered by traumatic life experiences (Kauer-Sant’ Anna *et al.* 2007) as well as by acute and chronic stress in animals (Duman & Monteggia 2006). The fact that these molecules are altered as a consequence of stress does not preclude, of course, that they may also play a causal role in the development and/or progression of mental disorders (Kapczinski *et al.* 2008); however, it does suggest that they are likely to be nonspecific as diagnostic biomarkers.

Interestingly, alterations in both cytokines and cortisol levels have also been found in non-psychiatric medical conditions, such as coronary heart disease (Mendall *et al.* 1997; Danesh *et al.* 2004; Smith *et al.* 2005) and cancer (Martín *et al.* 1999; Rich *et al.* 2005). It is possible, thus, that low-grade inflammation related to psychiatric illness could be linked to the increased clinical morbidity and mortality observed in patients with mood disorders and schizophrenia, particularly due to cardiovascular causes (Olfson *et al.* 2015; Ösby *et al.* 2016). Importantly, systemic inflammation can also have repercussions in the brain, and could be part of a feedback loop that leads to persistence of symptoms in disorders such as major depression (Gold 2014; Wohleb *et al.* 2016).

Thus, although the word “biomarker” is frequently used in a diagnostic sense, most of the literature using the term seems to focus on nonspecific markers of general psychopathology. This is not to say that these markers are devoid of clinical application, as there are multiple potential applications for biomarkers besides diagnosis (Davis *et al.* 2014) – the case has been made, for example, to use BDNF and cytokine levels as markers of illness activity (Fernandes *et al.* 2013) or progression (Kapczinski *et al.* 2009). However, the fact that the literature on biomarkers is still very fragmented across DSM-defined disorders (only 11% of articles in our sample compared markers across more than one disorder, and this percentage has actually decreased over time) probably limits the understanding of these molecules in this sense, and seems to argue in favor of transdiagnostic approaches for their study (Cuthbert & Insel 2013; Lancet Psychiatry 2016).

Evaluation of the clinical utility of biomarkers is also limited by the types of articles that have been found to be most prevalent in the literature – namely, crosssectional studies focusing on comparisons between patients and healthy controls. Although these studies are obviously necessary as starting points for further research, and have built a solid case for the robustness of biomarker alterations in psychiatric disorders, they are probably not representative of the typical situation in which a marker would be clinically useful (Perlis 2011; Kapur *et al.* 2012). Adding to this, measures of diagnostic efficacy that quantify how much a biomarker adds to clinical reasoning, such as ROC curves or odds ratios, were very infrequent in our sample – in fact, they have been *less* reported for recent biomarkers than they were for the evaluation of the dexamethasone suppression test three decades ago (Arana 1985). Taking all of this into account, the available literature on peripheral markers, although extensive and convergent in many points, is still insufficient to adequately assess their clinical potential.

In a more qualitative note, studies varied widely in terms of methodology, sample size and quality of reporting. Particularly notable were discrepancies between the numbers of measured variables and/or proposed analyses in the methods sections and those reported as results. It was not infrequent for articles to examine a large number of biomarkers, correlate them with many clinical variables, and report only significant associations, in a practice best described as selective outcome reporting bias (Dwan *et al.* 2008). The combination of a large number of outcomes with selective reporting will inevitably increase the possibility that reported findings are false positives (Ioannidis 2005), especially in the absence of multiple comparison corrections, which were infrequently performed. Importantly, it also leads to nonreporting of negative findings, and thus bias attempts to systematically review the existing literature.

Meta-analysis of available data can solve some of these issues, and metaanalytic comparisons have shown variations in the assessed biomarkers to be generally robust in common psychiatric disorders (Table 3). However, meta-analyses are intrinsically limited by the quality of the data; thus, their results can be influenced by publication and outcome reporting biases, as recently studied for markers of bipolar disorder (Carvalho *et al.* 2016). Moreover, they have been most successful for comparisons that are prevalent among studies – i.e. those between patients and controls, and eventually between disease states or pre- and post-treatment levels.

Attempts to correlate biomarkers with symptom severity, illness progression or response to treatment have yielded less consistent findings in meta-analyses (with a few exceptions), probably because these correlations were not only less frequent, but also usually presented as secondary analyses, and thus more susceptible to bias.

Such a general overview of the literature naturally has its limitations. First and foremost, the use of the term “biomarker” in our searches was a deliberate choice for specificity over sensitivity, and probably led many studies referring to peripheral markers by other terms to be missed. Moreover, in some articles the markers we assessed were only one among many studied, and not necessarily the main focus of the work – thus, the prevalence of some analyses might have been larger if we had limited ourselves to studies focused on those markers. Finally, our choice to provide a bird’s eye view of the literature has led us to focus on the rule rather than on the exceptions – thus, one should keep in mind that there are several articles in our sample that do go beyond patient vs. control comparisons. Moreover, although in terms of prevalence they constitute a minority, the absolute number of these articles has increased as the biomarker literature grows as a whole.

In spite of these limitations, we believe that our general conclusion – namely, that the most frequently studied peripheral markers in psychiatry are nonspecific markers of psychopathology across multiple disorders – is probably correct. In this sense, it is useful to discuss ways in which the literature can change in order to better assess their clinical utility. First and foremost, it is important to define what one means by “biomarker” on a case-by-case basis, as there are many ways in which a molecule might behave as a marker (Davis *et al.* 2014). Moreover, we must move from studies chasing statistical differences between groups to those that evaluate whether a biomarker can make a difference in clinical decision processes (Perlis 2011; Kapur *et al.* 2012; Lancet Psychiatry 2016). Finally, these studies will likely require more rigid design and larger statistical power – in this sense, development of reporting guidelines for biomarker studies (Gnanapavan *et al.* 2014) pre-registration of study protocols (Kivimaki *et al.* 2013) and formation of multicenter consortia (Sullivan 2010) are foreseeable ways to improve reproducibility. Such changes might ultimately depend on reforming incentive systems for science, which currently favor translational promise over assessment of clinical utility (Joyner *et al.* 2016) and stimulate small-scale studies that maximize the possibility of significant results (Higginson & Munafò 2016; Smaldino & McElreath 2016).

If these changes indeed take place, peripheral biomarker research could follow the path of genetic epidemiology, in which underpowered studies with limited reproducibility (Siontis *et al.* 2010; Bosker *et al.* 2011) have gradually been replaced by much larger studies from multicenter consortia (Sullivan 2010) that can control for the number of comparisons performed and yield more reproducible data (Collins *et al.* 2013). Our data shows that this has not yet happened in the peripheral marker field: aside from weak trends for an increase in sample size and a decrease in the frequency of cross-disorder comparisons, hardly any of the literature patterns we studied seems to have changed significantly over a 20-year period. This picture could change in the near future, however, as the results of ongoing large-scale initiatives in biomarker research (e.g. Kennedy *et al.* 2012; Kessing 2016) become available.

An open question that arises from our data is whether the transdiagnostic nature of the markers we have studied is actually a general rule for peripherally measured molecules. It might be expected, after all, that serum markers will be limited in their correlations with symptoms produced by the highly specialized anatomy of brain circuits. Moreover, the fact that most single molecules have very modest impacts in the risk for psychiatric illness has been made clear by genomic studies (Kendler 2013), and is likely to hold true at the peripheral marker level. Thus, it is possible that serum markers in general will tend to be transdiagnostic, and that most molecules might prove to be no more specific than the ones we focused on. This could feasibly be improved by ‘omics’ approaches evaluating multiple markers at a time; however, at least in our sample, these articles were relatively uncommon, and although some studies have suggested that combinations of markers might hold greater diagnostic specificity (Domenici *et al.* 2010; Schwarz *et al.* 2012; Frye *et al.* 2015), such findings await independent replication.

Even if most peripheral markers prove to be nonspecific, however, biomarker research can still hold its value. Alterations in the markers we have studied seem to be generally reproducible across psychiatric disorders, suggesting that the consequences of chronic stress might represent a common pathway in the evolution of mental illness. Thus, research in this field might shed light on the connections between psychiatric and medical illness, yield insights on pathophysiology, and perhaps provide ways to assess risk, disease progression or severity, especially if various markers are used in concert (Kapczinski *et al.* 2009, 2010). It can also increase our knowledge of the consequences of chronic stress on the brain and body, and help in the creation of transdiagnostic approaches to bridge the gaps between psychiatric research and neuroscience, or between psychiatry and other fields of medicine (Cuthbert & Insel 2013). For this to happen, however, the field needs to taper down its initial optimism, acknowledge that most peripheral markers are not likely to be diagnostic or specific, increase the rigor of its approaches and focus on questions that can drive research from statistical to clinical significance.

## AUTHOR CONTRIBUTIONS

O.B.A. and J.V.P. designed the study. J.V.P. performed literature searches. T.C.M. programmed the text-mining algorithm. J.V.P., T.C.M. and O.B.A. extracted article data and cross-checked data for consistency. T.C.M., O.B.A. and J.V.P. designed figures and tables. O.B.A. wrote the initial version on the manuscript. All authors provided critical input in revisions of the manuscript.

## ACKNOWLEDGEMENTS

The authors are deeply thankful to Marcia Kauer-S ant’Anna and Flávio Kapczinski for thoughtful discussions and constructive criticism on previous versions of the manuscript.

## FINANCIAL SUPPORT

This work was supported by FAPERJ grant E-26/201.544/2014 to O.B.A. and by a CNPq scholarship to T.C.M.

## CONFLICT OF INTEREST

The authors declare no conflict of interest pertinent to the current work.

